# Measurement of skeletal muscle fiber contractility with high-speed traction microscopy

**DOI:** 10.1101/733451

**Authors:** M Rausch, D Böhringer, M Steinmann, DW Schubert, S Schrüfer, C Mark, B Fabry

## Abstract

We describe a technique for simultaneous quantification of the contractile forces and cytosolic calcium dynamics of muscle fibers embedded in three-dimensional biopolymer gels. We derive a scaling law for linear elastic matrices such as basement membrane extract hydrogels (Matrigel) that allows us to measure contractile force from the shape of the relaxed and contracted muscle cell and the Young’s modulus of the matrix, without further knowledge of the matrix deformations surrounding the cell and without performing computationally intensive inverse force reconstruction algorithms. We apply our method to isolated mouse flexor digitorum brevis (FDB) fibers that are embedded in 10 mg/ml Matrigel. Upon electrical stimulation, individual FDB fibers show twitch forces of 0.37 µN ± 0.15 µN and tetanic forces (100 Hz stimulation frequency) of 2.38 µN ± 0.71 µN, corresponding to a tension of 0.44 kPa ± 0.25 kPa and 2.53 kPa ± 1.17 kPa, respectively. Contractile forces of FDB fibers increase in response to caffeine and the troponin-calcium-stabilizer Tirasemtiv, similar to responses measured in whole muscle. From simultaneous high-speed measurements of cell length changes and cytosolic calcium concentration using confocal line scanning at a frequency of 2048 Hz, we show that twitch and tetanic force responses to electric pulses follow the low-pass filtered calcium signal. In summary, we present a technically simple high speed and high throughput method for measuring contractile forces and cytosolic calcium dynamics of single muscle fibers. We expect that our method will help to reduce preparation time, costs, and the number of sacrificed animals needed for experiments such as drug testing.

**Statement of significance:** We describe a high speed, high throughput method for the simultaneous measurement of contractile force and cytoplasmic calcium dynamics following electrical pulse stimulation of muscle fibers embedded in a 3-dimensional biopolymer matrix. In contrast to the classical approach of attaching muscle fibers to a force-transducer, our method allows for a highly efficient, parallel analysis of large numbers of fibers under different treatment conditions.

## Introduction

The measurement of contractile forces is of central importance to assess the function of skeletal muscle tissue in human and animal models, and for quantifying the effect of drugs to treat genetic or age-related muscle diseases. Skeletal muscle cells, however, lose a large part of their initial contractility during prolonged cell culture (1). The contractile behavior of bioartificial muscles tissue cultured from myoblasts or induced pluripotent stem cells differs from freshly isolated muscle tissue regarding, for example, maximum contractility, calcium release and uptake kinetics, the expression levels of ion pumps, or the arrangement of RYR1 receptors (2). Therefore, the standard approach to perform such measurements is to mount either whole muscle explants or freshly isolated single muscle fibers between a force transducer and a length driver (3, 4).

A single muscle explant such as the flexor digitorum brevis (FDB) muscle can contain several hundred individual fibers, but their isolation and attachment to miniaturized mechanical force transducers and length drivers is exceedingly laborious and time-consuming, and this approach is therefore not suited for testing a large number of fibers (5–7). While force measurement in isolated whole muscle preparations (e.g. the extensor digitorum longus (EDL)) or soleus of mice or rats (8) is technically less demanding, there are ethical considerations that impose limits on the number of measurements. Moreover, the diffusion of substances from the organ bath into the muscle is hindered by the fascia and muscle tissue itself so that primarily the response of the outer layer of muscle fibers is measured.

In the present study, we adapt the method of traction force microscopy to measure the contractile response of isolated FDB fibers to electrical stimulation. The principal idea of traction force microscopy is to attach cells to an elastic matrix and to measure the matrix deformation in response to changes in contractile force, e.g. during electric field stimulation. If the elastic matrix properties are known, the traction forces that the cells exert onto the matrix can be computed from the measured matrix deformations.

For the case where the cells are attached to the surface of a flat, linearly elastic matrix (2-dimensional traction force microscopy), it is sufficient to measure the matrix deformations only at the matrix surface. Computationally and experimentally efficient methods for high-throughput measurements with high temporal resolution have been developed for this purpose (9, 10). However, freshly isolated muscle fibers do not attach well to such matrix surfaces and must therefore be embedded in a 3-D matrix. Computational methods for traction force reconstruction in 3-D are considerably slower, however, and the measurement of 3-D matrix deformations e.g. with confocal microscopy, is also time consuming (11) and therefore unsuitable for situations where the contractile forces rapidly change over time, such as during electrical pacing.

Here, we present a method to measure contractile forces of matrix-embedded muscle fibers that is both computationally efficient and experimentally simple and fast so that contractile responses during electric field stimulation can be measured at arbitrarily high frequencies in real-time. Our method exploits the approximately cylindrical symmetry of skeletal muscle fibers so that total muscle contractility can be calculated from the cell shape, contractile shortening and matrix stiffness with a simple scaling equation. In parallel to the measurement of contraction, we used the organic calcium indicator Mag-Fluo-4 to quantify calcium release and reuptake by the sarcoplasmic reticulum (SR).

## Material and methods

### Isolation of FDB fibers

All animal work was conducted according to national and international guidelines and approved by the cantonal veterinary services Basel Stadt. Flexor digitorum brevis (FDB) muscle is dissected from adult 6-9 weeks old male C57BL/6 mice, which are killed by decapitation after anaesthetizing them with isoflurane (4% in air). The FDB muscle is enzymatically dissociated for one hour at 37 °C and 5% CO_2_ in Tyrode’s buffer (138 mM NaCl, 2 mM CaCl 2, 1mM Mg Acetate, 4 mM KCl, 5 mM Glucose, 10 mM HEPES, pH 7.4) containing 2.2 mg/ml collagenase I (Sigma). During this incubation period, 35 mm diameter cell culture dishes with a 22 mm diameter glass bottom (Ibidi, Munich, Germany) are prepared by pipetting and evenly spreading 50 µl of 10 mg/ml Matrigel (Corning) onto the glass window and transferring the dish to an incubator at 37°C, 5% CO_2_ and 95% humidity to initiate the polymerization of the approximately 130 µm thick Matrigel film. To track the contraction-induced deformation field of the Matrigel surrounding the cells during experiments (which we use here only for the verification of the method and which is therefore not needed for routine measurements of contractile forces), we mix 7 µm diameter fluorescent latex beads (MF-FluoGreen-Particles, Particles GmbH, Berlin, Germany) to the Matrigel solution prior to polymerization.

After enzymatic muscle dissociation for one hour, 150-200 intact muscle fibers are manually isolated using fire polished pipette tips, mixed with 100 µl of 10 mg/ml ice-cold Matrigel, pipetted onto the previously polymerized Matrigel film in the center of the glass bottom dishes, and polymerized in an incubator at 37°C, 5% CO_2_, 95% humidity for 30 min. Because the polymerization process starts only after the Matrigel has sufficiently warmed, the cells have time to sink to the bottom of the previously polymerized Matrigel film and align horizontally, which is important for high-speed force measurement as it ensures that the cell shape can be captured from images obtained at a single focal plane. Finally, the dishes are covered with 3 ml Dulbecco’s Modified Eagle Medium (DMEM) supplemented with 10% FCS and 1% Penicillin-Streptomycin and kept in the incubator at 37°C until imaging is started. For measurements of cytosolic calcium concentration, FDB fibers are stained at room temperature for 20 min prior to imaging with 5 µM of the calcium-sensitive dye mag-fluo-4 AM (KD ~22 µM, Invitrogen) dissolved in Tyrode’s buffer.

### Microscopy

Measurements are carried out at room temperature using an inverted Olympus FV3000 confocal microscope equipped with a 20x (NA 0.75, air) objective and a resonance scan head for fast imaging. Two different imaging protocols are used:

a. For analyzing fiber contraction and bead displacements, images from the green fluorescent channel (ex488nm/em500-600nm) and the transmitted light channel are recorded in parallel at 30 frames/s and a spatial resolution of 512 x 512 pixels corresponding to 636 x 636 µm. 160 images are recorded in total for each stimulation cycle. The minimum time between two consecutive stimulation cycles is 8 sec. To record the data from all fibers in a dish as quickly as possible, a low-resolution map of the whole dish is recorded first for efficient navigation and selection of intact fibers for imaging.
b. For simultaneous calcium and force measurements, single line scans at a wavelength of 488 nm and a scan frequency of 2048 Hz (bi-directional scanning mode) are performed along a fiber. Intracellular calcium is obtained from the emitted light scan profile (500-600 nm) averaged over the length of the fiber. Before measurements, a single bright-field image of the relaxed muscle fiber is recorded to obtain the cell geometry.

### Electrical pulse stimulation

A high current stimulator (custom-made by SI Heidelberg, Germany) in combination with two platinum wire electrodes placed 10 mm apart is used for electric pulse stimulation. The amplitude of the bipolar pulses with a rectangular waveform is ± 9 V and the pulse duration is 1 ms. We use either single pulses delivered > 8 s apart for measurement of twitch forces, or pulse trains of 300 ms total duration with pulse frequencies of 10, 25, 50, 75 and 100 Hz. The timing of image acquisition and electric pulse stimulation is controlled by LabChart software and PowerLab data acquisition hardware (ADInstruments, Australia).

### Rheological Measurement

A cone-plate shear-rheometer (Discovery HR-3, TA Instruments) is used to measure the material properties of the 10 mg/ml Matrigel hydrogels. A 850 µL sample is polymerized at 37°C in a 65µm gap between plate and cone (diameter 40 mm, cone angle 2°). We measure the shear modulus for various strains (0.05 % −3 %) at a fixed frequency of 0.1 rad/s. We then measure the shear modulus for various frequencies (0.1 rad/s −10 rad/s) at a fixed strain of 0.5%. With a further amplitude sweep (from 0.05 % −100%) we measure the shear modulus at high strains to test for non-linear behavior (Fig. S2). The Young’s modulus is calculated from the storage modulus measured at 1 Hz assuming a Poisson ratio of 0.25 (11). The resulting value of 202.2 Pa ± 7.9 Pa is in agreement with previously reported measurements of Matrigel (12).

### Finite-element representation of a muscle fiber

We model the geometry of the experimental setup as a sphere of Matrigel with 2 mm radius (Fig. S1a). From confocal images, we determine the length and width of a muscle fiber in its relaxed state and place a cylindrical inclusion of the same length and diameter as the cell in the center of the sphere (Fig. S1b). The volume of the sphere outside the cylindrical inclusion is then filled with tetrahedral elements using the open-source mesh generator *Gmsh* (13). To ensure numerical accuracy of the finite element computation, the local node density is chosen to increase linearly with the curvature of the geometry. Thus, the size of the tetrahedral elements increases from ~1 µm near the cylinder surface representing the cell surface to ~200 µm at the outer boundary of the sphere.

### Force reconstruction

To compute cell contractility and the material deformations induced by a contracting muscle fiber, we assign the appropriate material properties and boundary conditions to all elements of the tetrahedral mesh. Matrigel at a concentration of 10 mg/ml shows predominantly linear behavior under shear, compression and elongation, with a constant (strain-independent) Young’s modulus of 202.2 Pa (Fig. S2) and a Poisson ratio of 0.25, as measured with a cone-plate rheometer (Fig. S2), and with an extensional rheometer and with stretch experiments as previously published in (11).

For assigning appropriate boundary conditions, we obtain the uniaxial strain *ϵ* along the long axis of the fiber (x-axis) from the fiber length in the contracted (*l*_*con*_) and relaxed (*l*_*rel*_) state according to ϵ = (*l*_*rel*_ − *l*_*con*_)/*l*_*rel*_. We assume this strain to be constant over the entire muscle length. We furthermore assume symmetry (displacements at the cell center are zero and both cell ends contract by the same distance towards the center) so that the constant strain sets the axial displacements *d*_*x*_ of the finite element nodes along the cell surface according to *d*_*x*_ = 0.5 *ϵx*, whereby the origin of the x-coordinate is at the cell center. The radial displacements along the cylinder surface are assumed to be constant along the length of the cylinder and are computed assuming volume conservation (i. e. the cytoplasm is assumed to be incompressible). We allow forces to occur only at the element nodes at the cell surface and set them to zero elsewhere. Finally, we assume zero deformations at the outer boundary of the simulated volume, far away from the muscle fiber. We then estimate the deformation field surrounding the muscle fiber by iteratively minimizing the strain energy within the simulated volume using the finite element software SAENO (Semi-affine Elastic Network Optimizer, Steinwachs et al. 2016).

For each node at the cell surface, we thus obtain the traction force that causes the local surface displacements. For each node within the material, we obtain the deformation. For each node at the outer surface of the sphere, we obtain the residual tractions needed to ensure zero displacements. We determine the total contractility of the muscle fiber by summing up the x-components of all forces on the cell surface from one end of the cell to its center.

### Validation of FE-Simulations

To validate the simulations, we manually measure the displacements of fiducial markers in the gel surrounding the muscle fiber (Fig. 2) between the relaxed and contracted state of the fiber using the image annotation software Clickpoints (14). To quantify differences in magnitude and direction between predicted and measured matrix deformations, we calculate the Pearson correlation coefficient between the measured and predicted bead displacements in both x-and y-direction. To minimize the influence of measurement noise, we limit this analysis to fibers measured with 100 Hz tetanic stimulation, for which the largest matrix deformations are observed.

## Results

### Deformation field

During electric field stimulation with single pulses, the muscle cells contract for a duration of ~100 ms with maximum strains of approximately 4-5%. In most cases (exceptions are discussed in the following paragraph), fluorescent beads at both ends of the muscle fibers move in the direction of the cell’s long axis towards the muscle center by a similar distance as the muscle ends, confirming our assumption of a no-slip boundary condition between the cell and the matrix.

The magnitude of matrix deformations decreases with increasing distance from the cell surface. Movements of beads near the cell poles point towards the cell center, while beads near the cell center point away from the cell. This observation agrees qualitatively with our prediction from finite element simulations (Fig. 1a). Moreover, we find good quantitative agreement between measured and predicted matrix deformation vectors (Fig. 2) with a Pearson correlation coefficient of 0.9 or higher for most cells (Fig. S3, S4). However, in some cases, we find zero or close-to zero matrix displacements near the cell despite large cell contractions, indicative of poor adhesion between the cell and the matrix (Fig. S3f,g, S4f,g).

**Fig. 1:**
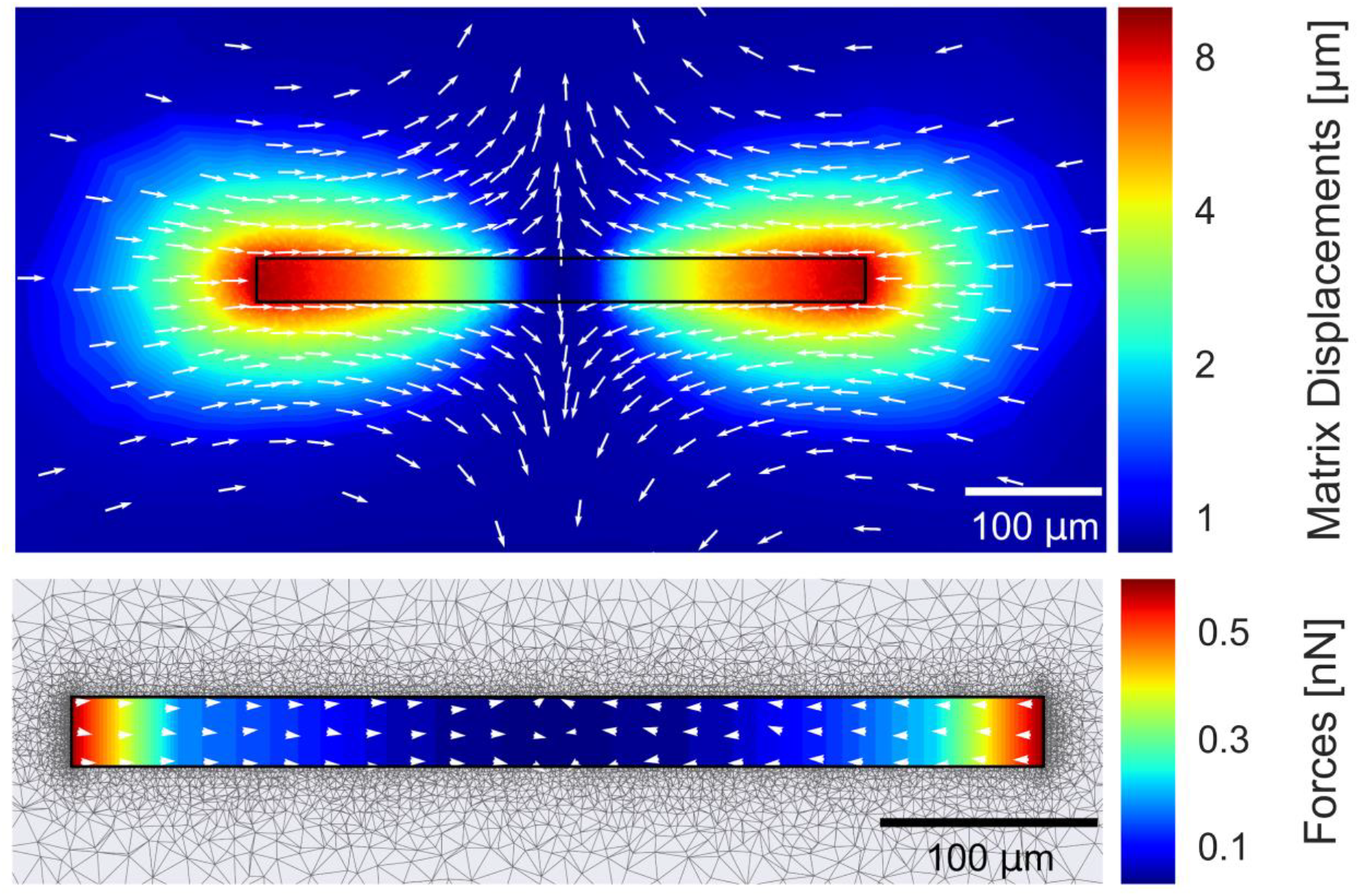
**a:** Reconstructed matrix deformations for a single contracting FDB fiber with a length of 446 µm, a width of 32 µm and 4% strain. A cross-section through the z-plane is shown. Displacement magnitude and direction are indicated according to colorbar and arrows. **b**: Reconstructed forces on the fiber surface. Absolute forces decrease towards the center. Total contractility is determined from the forces in x-direction summed over all surface nodes from one end of the fiber to its center. The force direction is predominantly aligned along the fiber axis as indicated by arrows. An outward normal directed force component due to volume conservation becomes visible for small displacements near the center.

**Fig. 2:**
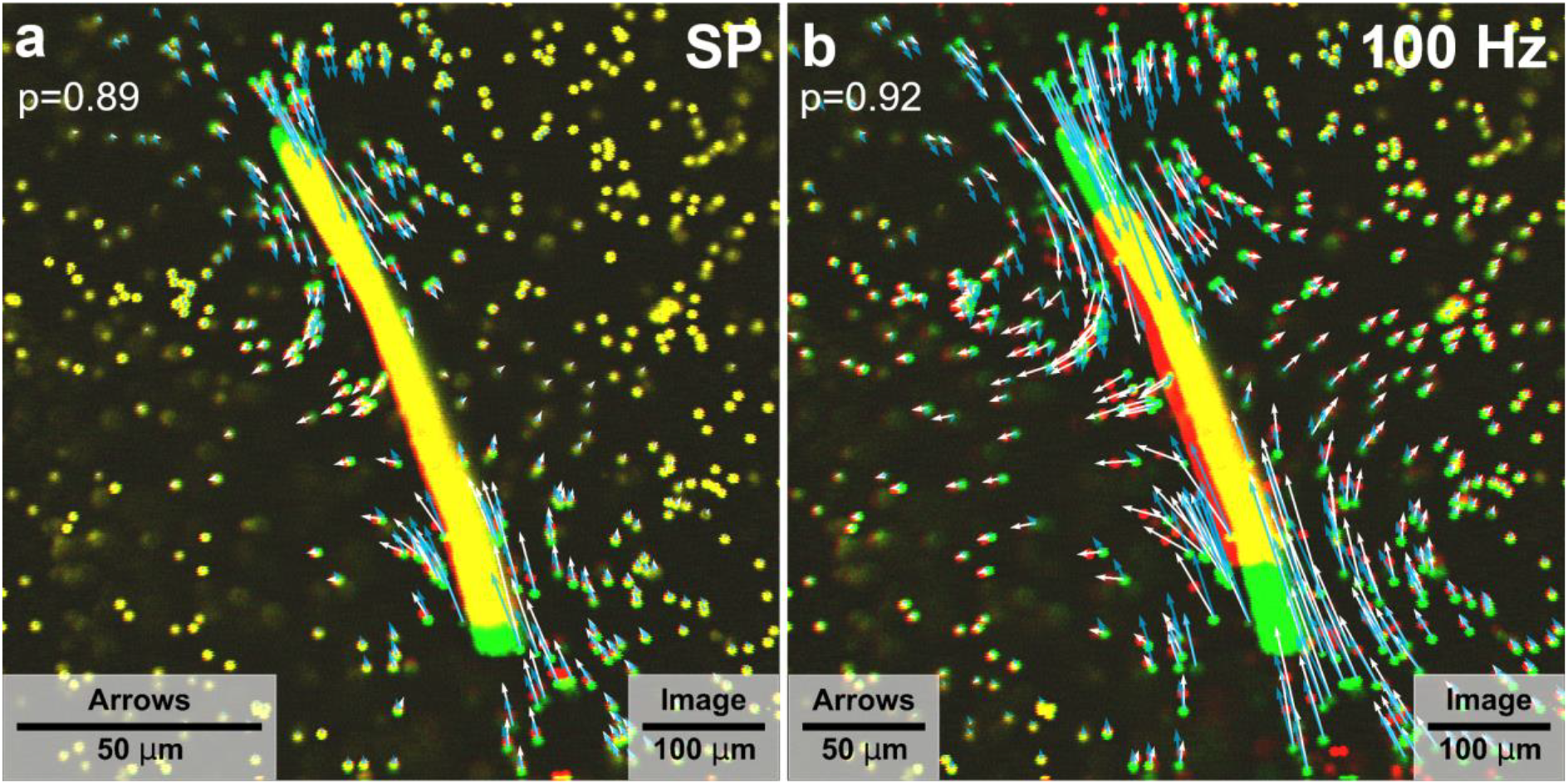
Matrix and fiber displacements in the mid-plane of the FDB muscle fiber. The relaxed state in green is superimposed with the contracted state in red. The FDB fiber is electrically stimulated with a single pulse (**a**) and a 100 Hz tetanic pulse train (**b**). White arrows indicate the measured matrix displacements, blue arrows indicate the predicted matrix displacement from FE modeling.

Unlike other traction microscopy methods where the measured 3-D deformation field serves as an input for the traction reconstruction algorithm, our method requires only knowledge of the elastic matrix properties, the cell length and cell diameter in the relaxed state, and the cell shortening during contraction. Therefore, the agreement between the predicted and the measured deformation field can be used as a validation of our method and justifies our highly simplified assumptions of cylindrical cell shape, constant cell strain and constant cell volume. Appreciable deviations between the measured and predicted deformation field arise predominantly when the cell is curved along its long axis or when the fiber is weakly attached to the surrounding matrix (Fig. S3, S4).

### Traction forces

During the process of iteratively minimizing the strain energy within the simulated matrix volume, the finite element network optimizer assigns traction forces to the surface of the cylindrical inclusion that account for the prescribed surface deformations and ensures that the boundary conditions are met. Upon convergence, we find the largest tractions at the two end faces of the cylinder representing the cell, whereas tractions at the side of the cylinder are small and moreover decrease towards the center. The direction of the traction vectors is predominantly aligned with the cell axis; a small normal force component arises for tractions near the cell center, due to the assumption of conserved cell volume (Fig. 1b). We compute total cell contractility by summing over all positive force vectors in x-direction.

### Scaling Law

For a series of simulations with different cell geometries, strains, and matrix properties, we calculate the deformation and traction force field. We find that the total contractility of muscle fibers scales linearly with matrix elasticity, linearly with fiber strain (Fig. 3a), quadratically with the fiber length (Fig. 3b), and linearly with fiber diameter plus a force offset at a vanishing diameter (long but thin cell) that corresponds to the force of a contracting line segment (Fig. 3c). The total contractility of a muscle fiber can thus be estimated from the fiber dimensions (diameter *d* and length *l* of the relaxed fiber), the fiber strain *ϵ* during contraction, and the Young’s modulus *E* of the Matrix according to

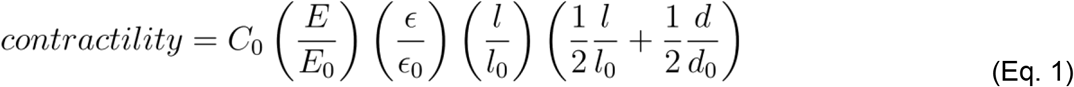

**Fig. 3:**
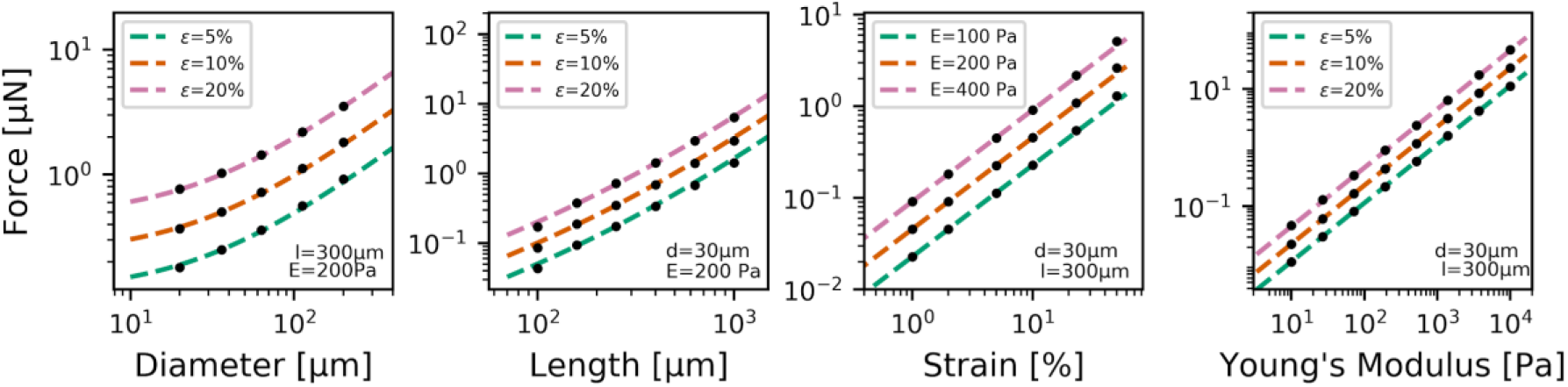
Cell contractility as a function of fiber diameter, fiber length, contractile strain and Young’s modulus of the surrounding matrix. Data points show the total contractility calculated from finite element analysis for differently shaped and differently contractile cells. Dotted lines are the predictions from Eq. 1.

Here, *C*_0_ = 0.455 µN denotes the contractility of typical reference fiber with length *l_0_* = 300 µm, diameter *d_0_* = 30 µm, and strain ε_0_ = 10%. *l* denotes the length, *d* denotes the diameter, and ε the strain of the muscle fiber to be investigated. *E* describes the Young’s modulus of the matrix, which is compared to a reference environment (10 mg/ml Matrigel) with *E_0_* = 200 Pa. Using this scaling law, the contractility of any fiber embedded in a linear elastic material can be evaluated using Eq. (1), without the need to perform additional finite element analyses or to measure matrix deformations. Moreover, we find excellent agreement between Eq. 1 and a full finite element analysis over a wide range of strains (1% −50%), cell lengths (100µm −1000 µm), cell diameters (20 µm −200 µm), matrix stiffness (10 Pa −10 kPa), and combinations thereof.

### Contractility Analysis

To demonstrate the practical applicability of our method, we compare contractility values computed with finite element simulations versus contractility values computed with Eq.1 in 14 cells from two independent fiber isolations (7 fibers each). The rms differences between both methods are 4 % (Fig. 4b), confirming the scaling law (Eq. 1). We find an average contractility of 0.37 µN ± 0.15 µN (Fig. 4a) when fibers are electrically stimulated by single bipolar pulses (±9 V bipolar pulses, 1 ms duration per pulse). The average contractilities from the two muscle isolations, when computed separately, are nearly identical (Fig. 4a), confirming the reproducibility of the method.

**Fig. 4:**
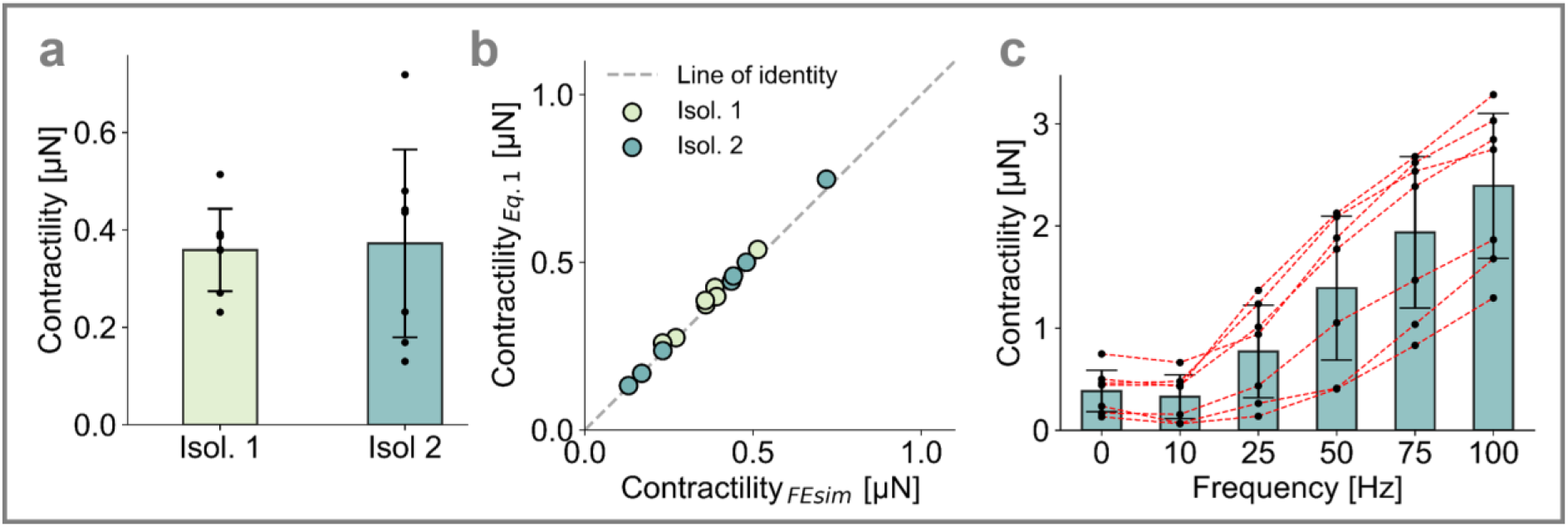
**a:** Contractile forces (0.37 µN ± 0.15 µN, mean ± sd) calculated using finite element simulations for two muscle isolations with 7 fibers each. Fibers are electrically stimulated with single pulses. Contractility values of individual fibers are shown by black markers. **b:** Single-pulse contractile forces calculated from cell shape changes using Eq. 1 versus forces calculated using finite element simulations. Each symbol corresponds to a single cell. Colors correspond to two different muscle isolations. **c:** Contractile forces (mean ± sd for 7 fibers of isolation #2), stimulated with single pulses (0 Hz) or 300 ms long pulse trains with pulse frequencies between 10 Hz −100 Hz. Forces are calculated using Eq.1. Black markers that are connected with red dashed lines indicate the response of individual cells.

### Contractile response to tetanic pulse sequences

When 300 ms long electrical pulse sequences are applied, the contractile force measured at the end of the sequence increases strongly with increasing pulse frequency between 10 Hz and 100 Hz (Fig. 4c). The contractility at 10 Hz is marginally smaller compared to single pulse forces. For a pulse frequency of 100 Hz, we observe the highest contractility of 2.38 µN ± 0.71 µN, corresponding to a tension of 2.53 kPa ± 1.17 kPa. Moreover, we find that cells with higher single-pulse contractility tend to display a higher contractility at other pulse frequencies.

### Drug testing

To demonstrate the applicability of the method for drug testing, we first treat FDB fibers with increasing doses of caffeine. Caffeine facilitates calcium release from the sarcoplasmic reticulum (15), which leads to higher muscle tension during electrical stimulation (16, 17). We confirm higher contractility of FDB fibers with increasing doses of caffeine both for single twitch and tetanus stimulation, except at a very high dose of 10 mM where the contractility during tetanic stimulation but not for single pulses begins to decline (Fig. 5a). This effect at high caffeine doses is not apparent from the muscle shortening data alone (inset Fig. 5a), which underscores that it is important to consider the whole cell geometry (length, diameter, strain) for estimating cell contractility.

**Fig. 5:**
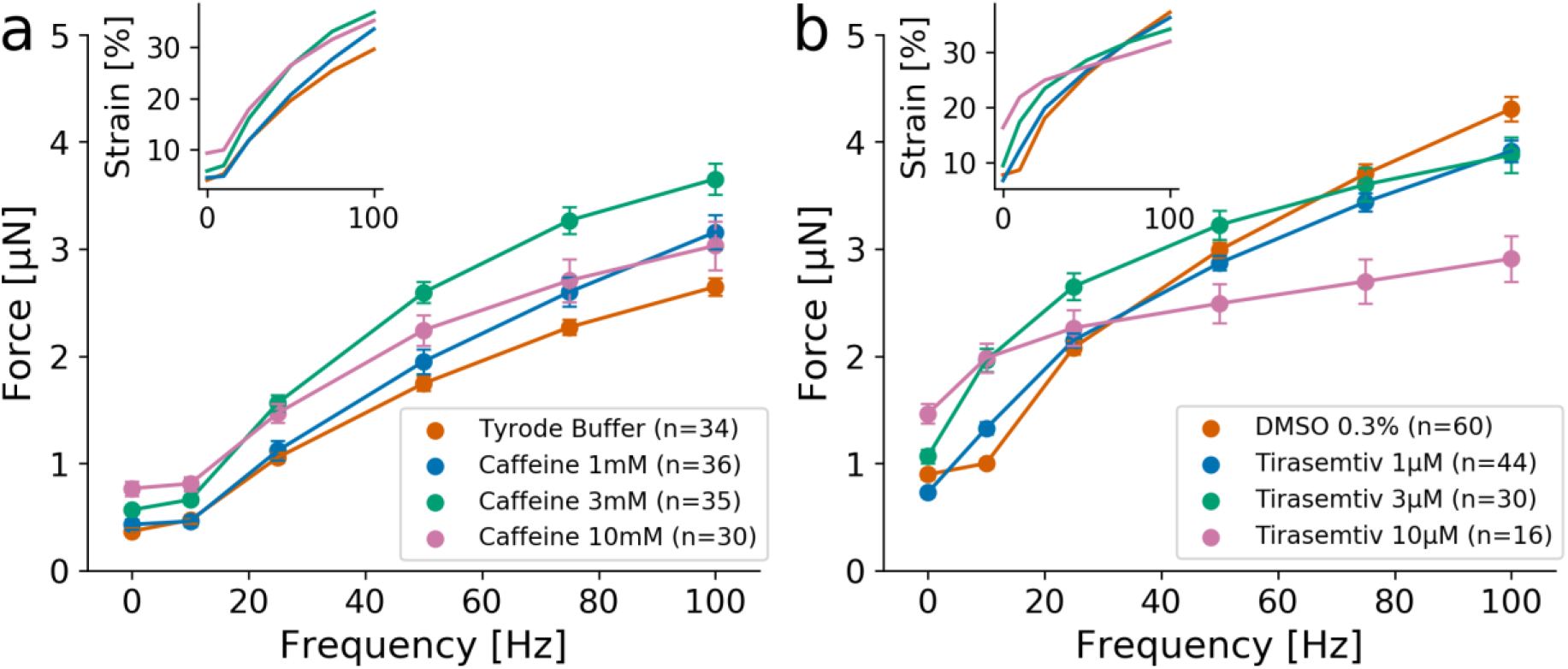
Force versus stimulation frequency relationship for FDB fibers treated with different concentrations of caffeine dissolved in Tyrode buffer (**a**) and Tirasemtiv dissolved in PBS with 0.3% DMSO (**b**). Insets show the fiber strain during contraction (mean of *n* FDB fibers as indicated in the legend). A dose dependent increase in force is observed for both drug treatments, except at the highest doses (10 mM for caffeine, 10 µM for Tirasemtiv) where the tetanus contractility decreases. Data points represent mean ± se of *n* FDB fibers as indicated in the legend.

Second, we treat FDB fibers with the drug Tirasemtiv (Cytokinetics Inc.), which is dissolved in DMSO (final DMSO concentration of 0.3 %). Tirasemtiv was developed as a muscle strength-increasing drug for the treatment of amyotrophic lateral sclerosis, a neurological disease that destroys motor neurons and leads to muscle degeneration. Tirasemtiv binds selectively to the fast skeletal troponin complex and slows the rate of calcium release from troponin C (18). Similar to caffeine-treated FDB fibers, we find an increase in contractility with increasing doses of Tirasemtiv especially for single pulses, but at higher frequencies, the effect of Tirasemtiv diminishes and ultimately reverses such that at stimulation frequencies of 75 Hz and 100 Hz, forces of drug-treated cells fall below those of DMSO control cells (Fig. 5b). Thus, a therapeutic effect of Tirasemtiv is only present at lower stimulation frequencies, in line with previous data (18).

### Simultaneous force and calcium measurement at high speed

We next combine force measurements with simultaneous calcium imaging using the fluorescent dye Mag-Fluo-4. From the single line scan signal of a confocal microscope (scanning frequency 2048 Hz, scan orientation aligned with the fiber), we extract both the average fluorescence intensity of the calcium dye and the muscle fiber length, which we convert to contractile force using Eq. 1 (Fig. 6).

**Fig. 6:**
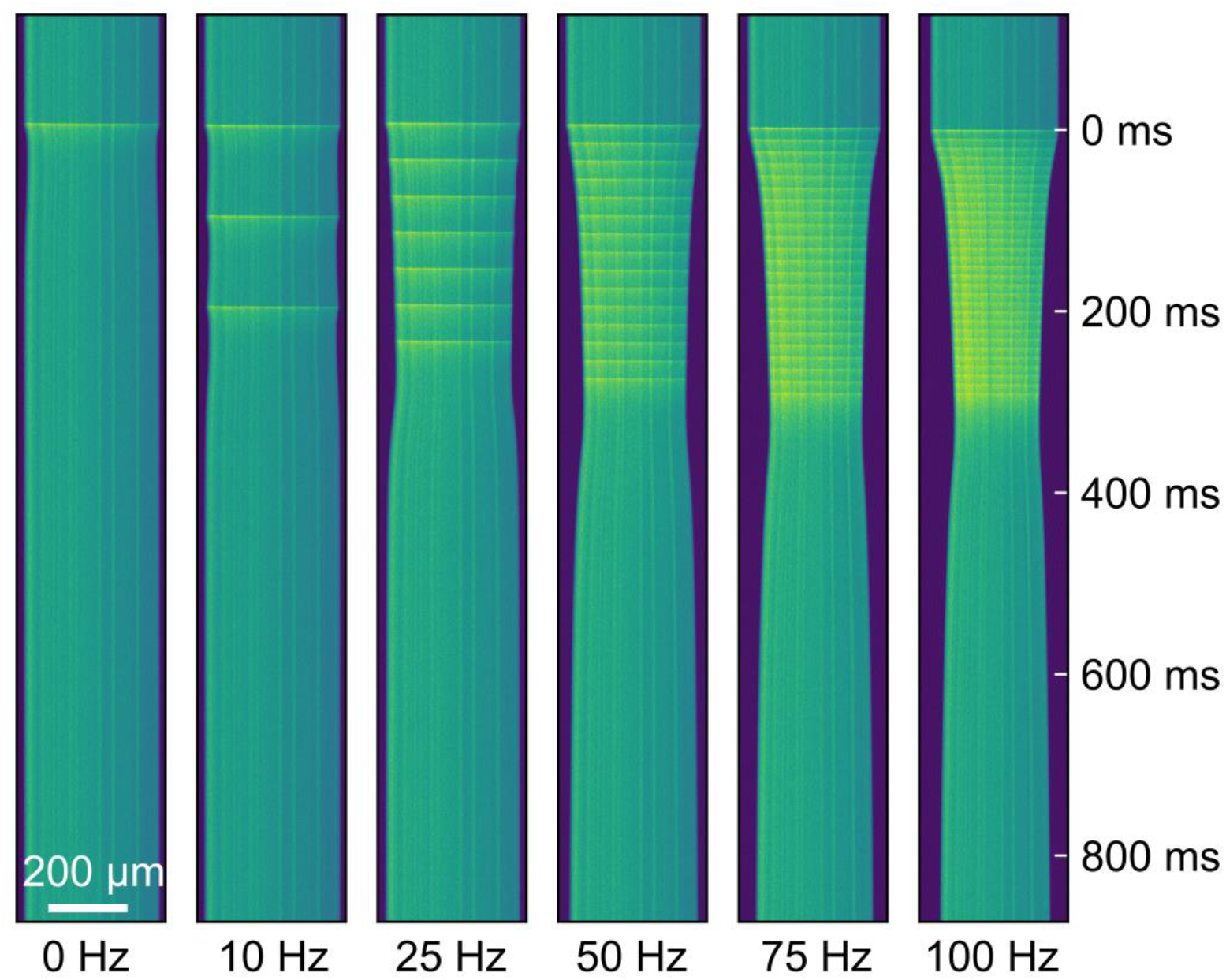
Kymograph derived from line-scan imaging demonstrating the increased Mag-Fluo-4 fluorescence and fiber shortening during electrical stimulation. Fluorescence intensity during electrical stimulation increases within few milliseconds (yellow-green horizontal bands). Shortening of the fiber is reflected by thinning of the green band, which corresponds to the length of the fiber.

We then measure the relationship between cytosolic calcium concentration and contractility for the same cell at different electrical stimulation frequencies (single pulse up to 100 Hz). We find that the maximum concentration of electrically induced calcium spikes remains nearly constant for different stimulation frequencies (blue line in Fig. 7). After application of a single electric pulse, the intracellular calcium concentration quickly peaks and then decreases within approximately 100 ms to baseline values. For stimulation frequencies above 10 Hz, however, the intracellular calcium concentration does not fully recover to baseline after each pulse and thus accumulates over time. We find that the contractile force (orange line in Fig. 7) follows approximately the low-pass filtered calcium signal (dashed line in Fig. 7) according to a first-order activation dynamics:

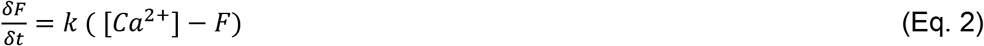

where *F* is the force (normalized to unity), [*Ca*^2+^] is the intracellular calcium concentration (normalized to unity), and *k* is the rate of contractile activation. In the case of FDB fibers, we find a rate of k = 10.28 s^−1^. The time constant of the low pass filter equals 96.7 ms.

**Fig. 7:**
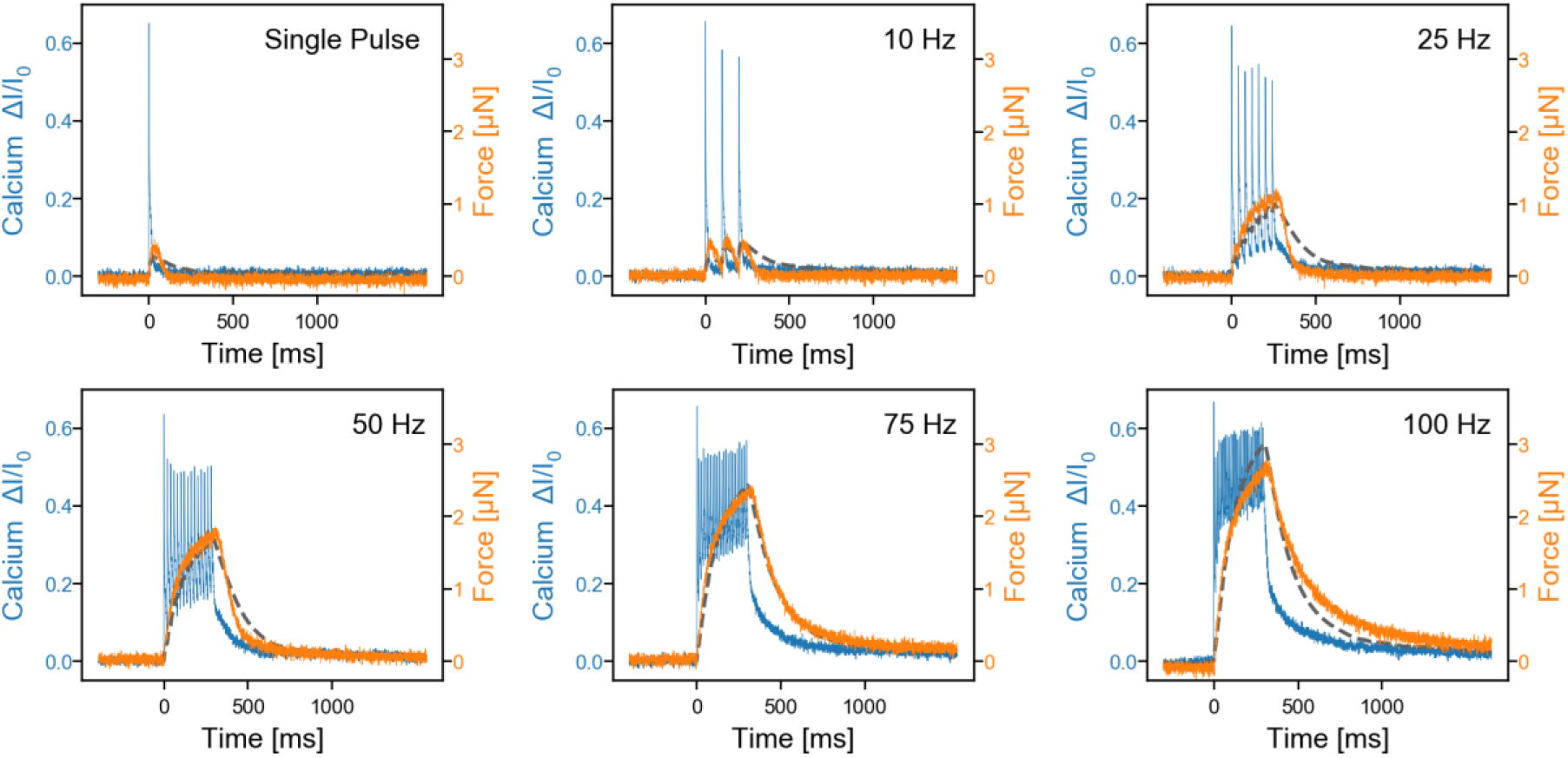
Simultaneous measurements of calcium changes (change in fluorescent intensity *Δ I* normalized to baseline intensity *I*_0_ measured in the relaxed fiber, blue line) and contractile force (orange line) for a single FDB fiber stimulated by electrical pulses of different frequencies. Contractile force can be predicted from the first-order resistor-capacitor (RC) low-pass filtered calcium signal (dark gray dashed line, time constant of the low pass filter = 96.7 ms), corresponding to a first-order activation dynamics according to Eq. 2.

## Discussion

We present a method for measuring contractile forces of single skeletal muscle fibers that are embedded in a 3-D extracellular matrix. By exploiting the approximately cylindrical geometry of isolated muscle fibers, our approach - unlike other traction microscopy methods - does not require knowledge of the 3-D matrix deformation field. Instead, only 4 input parameters are needed: the Young’s modulus of the matrix, the length and diameter of the relaxed muscle fiber, and the contractile strain. If the fibers are embedded in a linear elastic matrix such as Matrigel, contractile forces scale linearly with the Young’s modulus of the matrix and the contractile strain, quadratically with fiber length, and linearly with fiber diameter. Thus, for computing the contractile force of a given cell, it is not necessary to perform a finite element analysis; instead, the contractile force obtained from a single finite element analysis of a typical cell can be scaled with Eq. 1 to the geometry and contractile strain of any other cell. Cell geometry and contractile strain and therefore contractile force can be easily measured at high speed using bright-field or confocal microscopy.

We verify our method by comparing measured matrix deformations with matrix deformations predicted by finite element simulations. Since the measured matrix deformations do not serve as input values for our traction reconstruction algorithm, a good agreement between measured and predicted deformation vectors validates our choice of a highly simplified cylindrical cell geometry, no-slip boundary conditions at the cell surface, and our assumptions of a constant strain along the fiber axis and of cell volume conservation. Occasionally, however, we find substantial deviations between predicted and measured matrix deformations in cases where the cell contracts but adhere poorly to the matrix. Accordingly, matrix deformations even in close proximity to the cell surface are zero or near zero. To detect such cases and to ensure the quality of the measurements, it is highly advisable to add fiducial marker beads to the matrix.

A limitation of our method is that the resting muscle length (slack-length) cannot be controlled. After isolation, the fibers experience no restoring forces from connective tissue or antagonistic muscle tissue, and thus the length at which the fibers settle onto the surface of the pre-polymerized Matrigel surface is expected to be shorter than the optimal length that these fibers originally had in situ. At a resting length below the optimum length, which corresponds to a sarcomere spacing of approximately 2.4 µm for mouse FDB fibers (19), these fibers are expected to generate lower contractile forces (8).

Another limitation of our method is that the mechanical load against which the muscle fibers shorten during contraction can only be adjusted over a relatively narrow range. In particular, only weakly auxotonic contractions are possible with our setup, whereby the mechanical load is provided by the elastic restoring forces of the Matrigel. Compared to traditional measurements under isometric conditions, auxotonic contraction as employed in our setup makes the comparison between different measurements more complicated, as the mechanical load against which the muscle shortens depends on muscle geometry and matrix mechanical properties. Moreover, the stiffness of the Matrigel biopolymer can differ between batches and must be controlled for long-term studies or studies across laboratories. At a concentration of 10 mg/ml Matrigel, the hydrogel matrix has a Young’s modulus of 202.2 Pa (Fig. S2, Slater and Partridge 2017), which is much softer than the connective tissue of muscle with typical Young’s moduli on the order of 10 kPa (20). Thus, the low elastic load of the 10 mg/ml Matrigel matrix against which muscle fibers contract during electrical stimulation results in large strains of up to 40%.

According to the sliding filament theory, maximum tension decreases during muscle shortening, and most skeletal muscle fiber are unable to generate appreciable forces at contractile strains larger than 40% (4), which is close to the strain we see in FDB fibers undergoing tetanic stimulation. Sub-optimal resting length together with large contractile strains due to a very low elastic load of the matrix are therefore two likely reasons that may explain the relatively low contractile stresses of 2.53 kPa ± 1.17 kPa that we measure for 100 Hz tetanic stimulation. On the positive side, large contractile strains mean that the dynamic range of our method (from strains of 0.1% to 40%, corresponding to a dynamic range of more than two orders of magnitude) is sufficiently large to investigate compound effects (Fig. 5).

To increase the maximum tension of FDB fibers in our setup, hydrogel matrices with substantially higher Young’s moduli are required. These hydrogels, however, need to be functionalized with adhesive ligands that can support the transmission of the very large traction forces that fully differentiated skeletal muscle fibers are able to generate. In animals and humans, force is transmitted from the muscle to the tendon via the myotendinous junction (21). The myotendinous junction consists of membrane folds that form finger-like structures with a large surface between the sarcolemma and the collagen fibrils of the tendon. Specialized proteins such as the dystrophin-associated protein complex connect actin filaments and the cell membrane to the collagen fibrils and are able to transmit extremely high forces. In our setup, by contrast, transmembrane forces are transmitted by integrin-laminin-collagen complexes. In vivo, these complexes are important for withstanding stress during contractions but not for transmitting force, and therefore these connections are obviously much weaker compared to the myotendinous junction (22). In a series of pilot-experiments, we also tested other biopolymer gels (CELLINK, Boston, US), which after solidification by UV had a substantially higher stiffness compared to Matrigel (data not shown). However, FDB fibers detached from the matrix during contraction and did not return to baseline length, leaving a “tunnel” behind”.

Despite these limitations, data obtained with our method show good qualitative agreement with conventional isometric measurements obtained from intact FDB fibers. FDB muscle fibers from wildtype or transgenic mice have been previously used in numerous studies to investigate the effect of genetic mutations or aging on e.g. calcium dynamics, reactive oxygen production and mitochondrial function (23–25). FDB fibers are fully differentiated and maximally contractile already in newborn mice, and all secondary developmental processes such as the formation of the tethering structures for the sarcoplasmic reticulum and mitochondria are completed by the age of 8 weeks (26). In line with previously reported experiments on FDB fibers, we find a strong increase of contractile forces with increasing pulse frequencies of up to 100 Hz (27). Furthermore, the time course of contractile forces in response to a single electric pulse and pulse trains agrees with previous measurements (28). Also in agreement with previously reported data are our measurements of a dose-dependent increase in contractility after drug treatment with caffeine and Tirasemtiv (16, 18). Due to its simplicity and speed, our method to measure contractile forces can be easily combined with other high-speed imaging modalities such as Ca^2+^ measurements using fluorescent dyes. We find in FDB fibers that the contractile force closely follows the low-pass filtered calcium signal, which is consistent with the notion that muscle force is determined by a first-order activation dynamics according to Eq. 2, with a rate that is determined for example by the binding and unbinding kinetics of calcium to troponin. An alternative explanation, however, is given by the sliding filament model of Huxley (29). Accordingly, intracellular calcium leads to a nearly instantaneous activation of myosin motors, but the transmission of their combined forces to the extracellular matrix is suppressed due to frictional forces between myosin heads and actin binding sites as the thin and thick filaments slide against each other. The effect of internal friction on muscle force generation is described by Hill’s visco-elastic active-state model (3), which also takes the form as Eq. 2 if one neglects the parallel elastic element of the muscle, assumes that the internal force instantaneously follows intracellular calcium, and identifies the rate constant *k* as the total elasticity (sum of matrix elasticity and serial muscle elasticity) divided by the coefficient of friction from the viscous element of the muscle.

In summary, we have shown that our method can be used for drug testing, and that it can be applied to a range of questions in muscle physiology. The method does not require the laborious mounting of cells to micro-force transducers, and experiments of many cells can be conducted within the same cell culture dish, allowing for a massive increase in measurement throughput. Because several hundreds of muscle fibers can be isolated from a single muscle, our method greatly increases data yield and therefore may help reduce the number of sacrificed animals.

## Code Availability

The developed force reconstruction method is based on the network optimizer SAENO (11) and the finite element software Gmsh (13). The method is implemented in the python package *arnold*, which provides an interface to the finite element model, the simulations and the force reconstruction. The software is open source (under the MIT License) and accessible on GitHub (*https://github.com/davidbhr/arnold*). The figures in the work have been created using the Python packages Matplotlib (30) and Pylustrator (31).

## Acknowledgements

This work was funded by grants from the German Science Foundation (FA 336-12/1, TRR-SFB 225 projects A01 and A07), the Muscle Research Center Erlangen, the National Institutes of Health (HL120839) and the Emerging Fields Initiative of the University of Erlangen-Nuremberg (project “Novel Biopolymer Hydrogels for Understanding Complex Tissue Biomechanics”).

## Author Contributions

MR and MS developed the experimental technique, performed the cell isolations and all cell measurements. DS, DB and SS conducted the rheological measurements. BF, DB and CM developed the method for force measurements and analyzed the data. MR, BF, DB and CM wrote the manuscript. MR and DB contributed equally.

## Appendix

**Supplementary Fig. 1:**
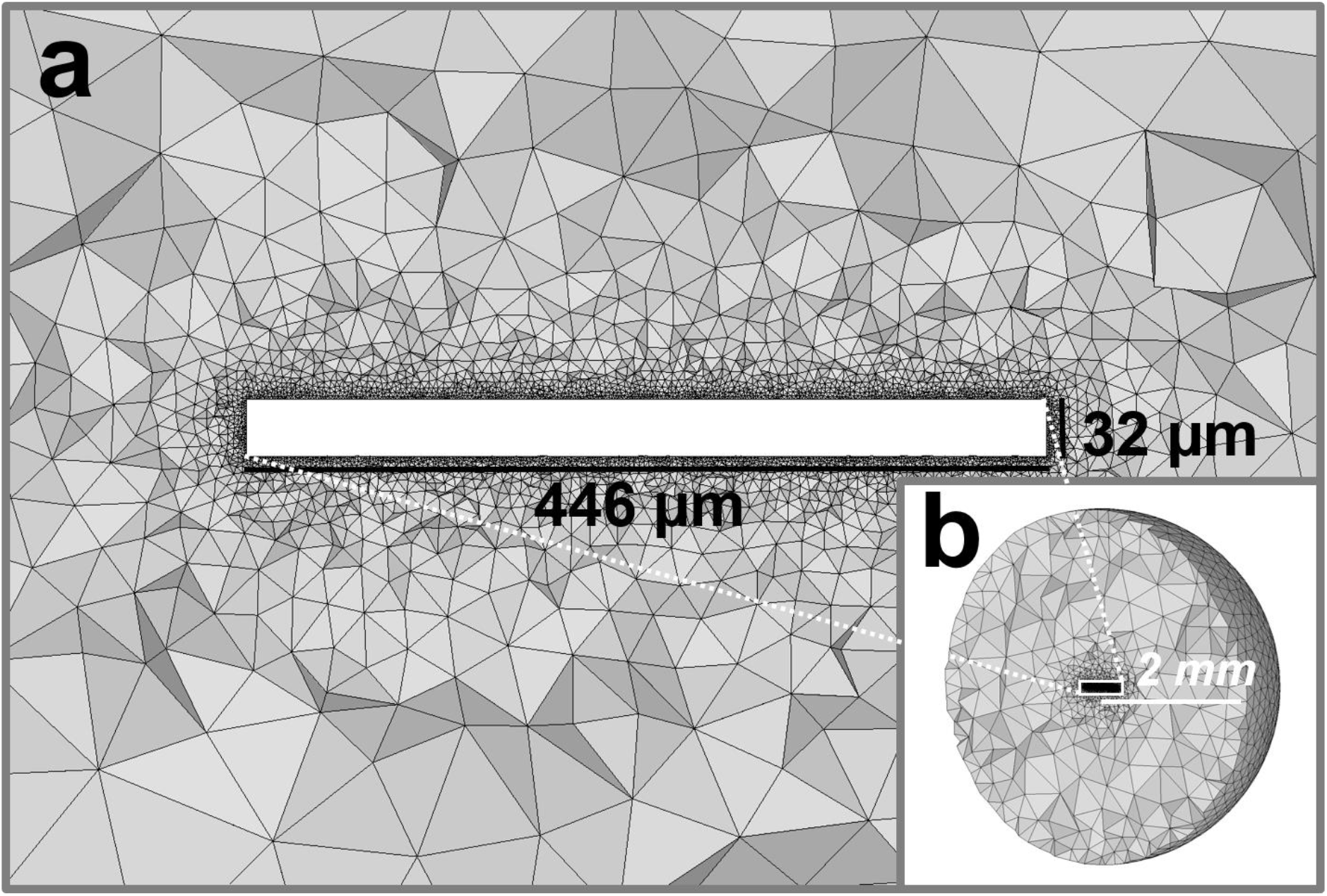
Tetrahedral mesh used in the finite element simulation: **a:** The cell is modelled as a cylindrical inclusion with a length of 446 µm and a diameter of 32 µm. **b:** The geometry of the experimental setup is modelled as a Matrigel sphere with a radius of 2 mm with the cell at its center.

**Supplementary Fig. 2:**
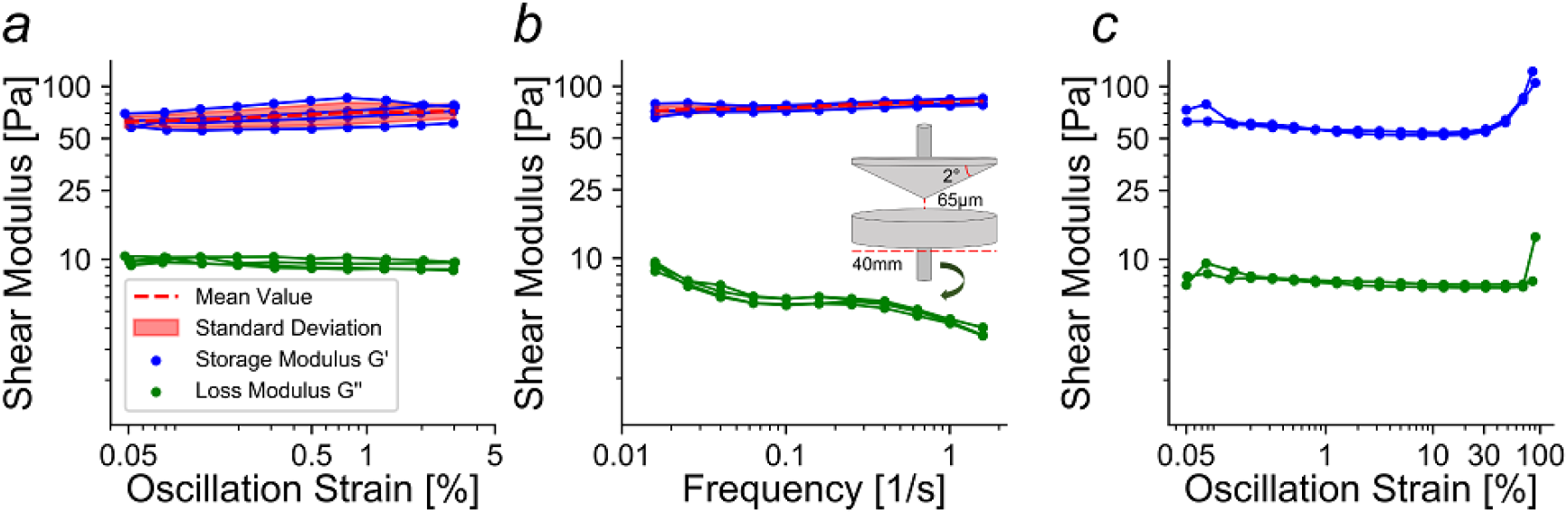
Cone-plate shear-rheometer measurements of 10 mg/ml Matrigel hydrogels. **a:** Shear modulus for various strains (0.05 % −3 %) at a fixed frequency of 0.1 rad/s. **b:** Shear modulus measured at a range of frequencies (from 0.1 −10 rad/s) at a fixed strain of 0.5%. **c:** Shear modulus at high strains (0.05% −100%). We observe predominantly linear elastic properties in all measurements.

**Supplementary Fig. 3:**
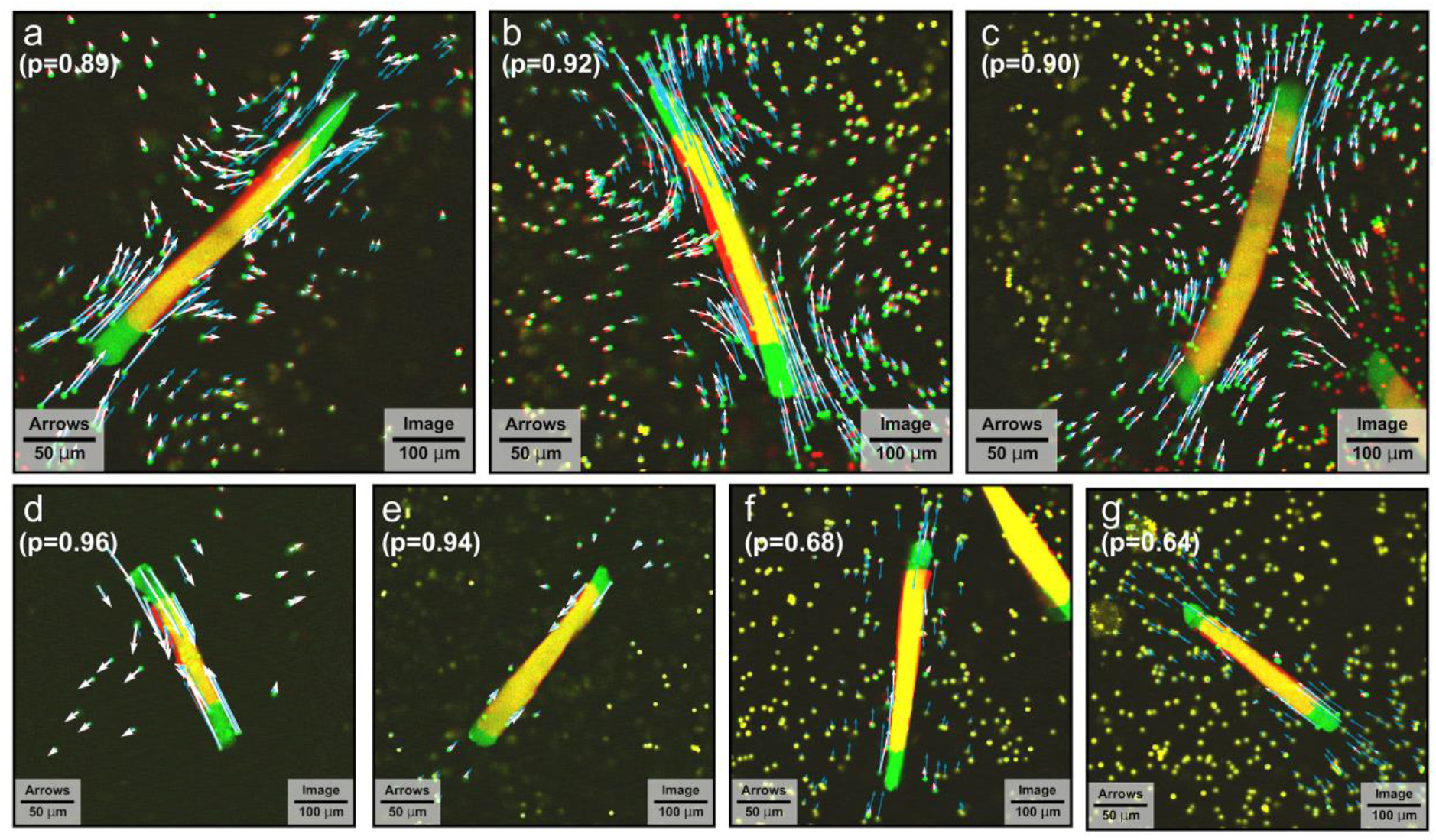
Matrix and fiber deformations in the mid-plane of the FDB muscle fiber. The relaxed state in green is superimposed with the contracted state in red. FDB fibers are electrically stimulated with 1 ms long pulses every 10 ms (100 Hz) for a duration of 300 ms. White arrows show the measured deformations, blue arrows show the predicted deformations from FE modelling. The correlation coefficient between predicted and measured deformations is 0.85 ± 0.12 (mean ± sd). Fibers in a-c show good agreement between predicted and measured deformations. In d) and e), the bead density is low, but the measured deformations are in good agreement with FE simulations. In f) and g) we find substantial deviations between predicted and measured matrix deformations. Small or absent bead movements indicate poor attachment between the FDB fibers and the surrounding matrix.

**Supplementary Fig. 4:**
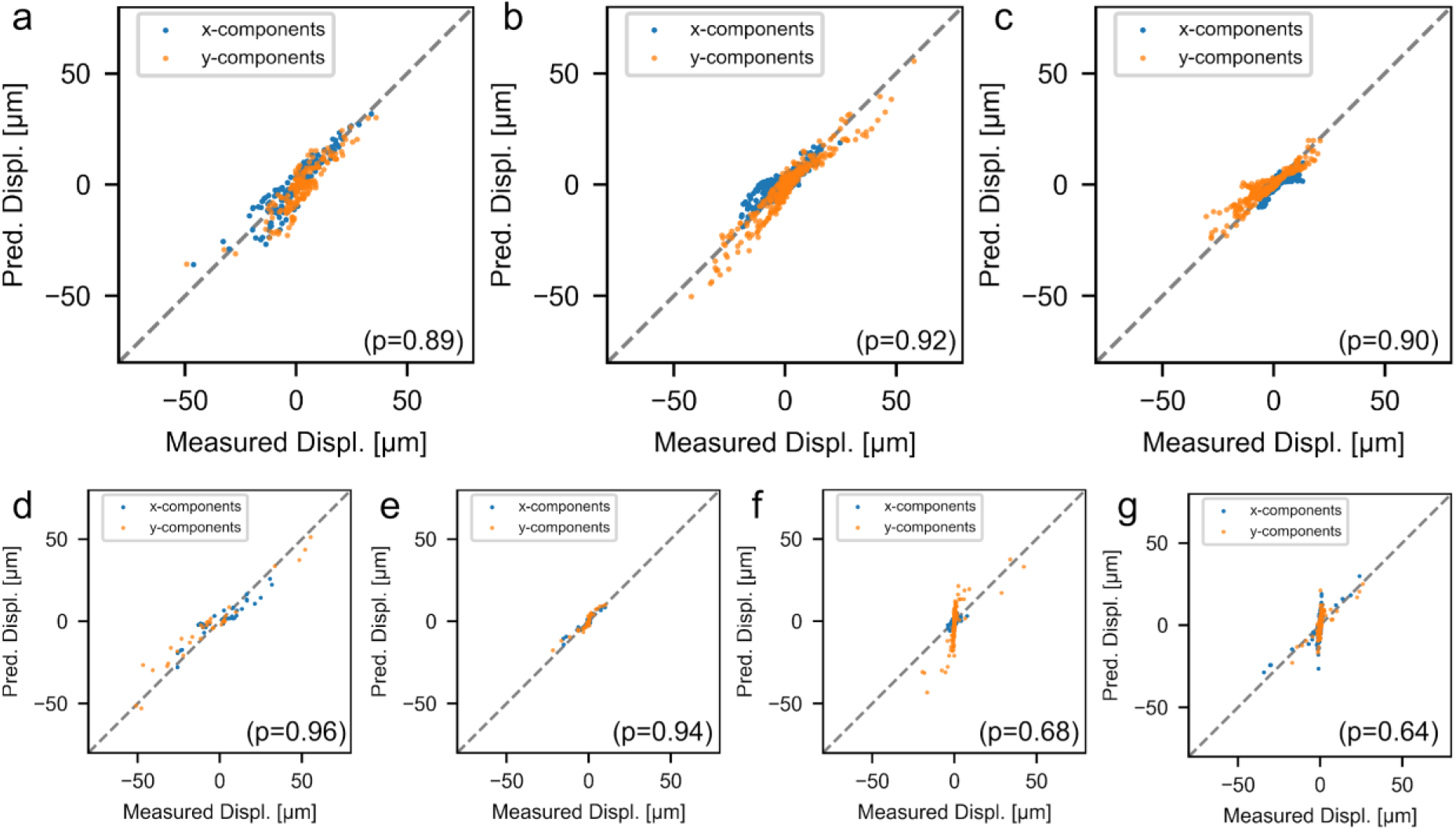
Predicted matrix deformations vs. measured matrix deformations in x-direction (blue) and y-direction (orange), for all FDB fibers shown in Fig. S3. Line of identity is shown in grey.

